# Degradation of intrinsically disordered proteins using proteolysis targeting nanobody conjugate

**DOI:** 10.1101/2024.11.27.622182

**Authors:** Xiaofeng Sun, Donglian Wu, Chengjian Zhou, Simin Xia, Xi Chen

## Abstract

Intrinsically disordered proteins (IDPs) play crucial roles in diverse cellular processes and are implicated in numerous diseases. However, due to the inherently unstructured nature, it is challenging to modulate their function using small molecule degraders or inhibitors. To address this, we introduced proteolysis targeting nanobody conjugate (PROTNC) composed of a nanobody that binds to an IDP, an E3 ligase ligand, and a cyclic cell-penetrating peptide for efficient intracellular delivery. We first showed a general-purpose PROTNC system capable to degrade different endogenous IDPs based on a co-condensate formation mechanism. We also introduced a straightforward strategy to degrade individual IDPs using specific PROTNCs equipped with a nanobody recognizing that particular IDP. Notably, we demonstrated the degradation of TPX2 which is an undruggable oncogenic IDP responsible for microtubule nucleation. This targeted degradation effectively inhibited cancer cell proliferation by disrupting bipolar mitotic spindle assembly in metaphase and directed cells to apoptotic pathway.

## Introduction

Intrinsically disordered proteins (IDPs), which make up nearly half of the human proteome play crucial roles in various cellular processes^1, 2^ and are associated with a range of human diseases like cancer and neurodegenerative disorders^3^. IDPs lack a fixed 3D structure under normal physiological conditions due to the presence of intrinsically disordered regions (IDRs) that exist as highly dynamic conformational states^4^. This characteristic makes IDPs challenging drug targets for classical structure-based drug design pipeline^5^ and other approaches^6^. Antibodies are another rapidly expanding category of drugs^7^, but they primarily bind cell surface receptors, making them difficult to interact with the wide range of IDPs found inside cells. Therefore, there is a high demand for general and innovative strategies to modulate the functions of IDPs, such as through direct alternation of the protein level via targeted protein degradation (TPD)^8^.

Proteolysis targeting chimeras (PROTACs)^9^ and alternative TPDs such as LYTACs^10^, Trim-Away^11^, ATTEC^12^, and ATNC^13^ are gaining wide attention in the fields of biology and medicine. These approaches utilize the ubiquitin-proteasome system (UPS), lysosome-endocytosis, or autophagy pathways to clear proteins^14^, making them potential strategies to deplete IDPs. However, traditional TPDs are bivalent small molecule dimerizers that consist of a ligand for recruiting a target protein and another module for directing to a degradation pathway^15^. For instance, a PROTAC typically carries a small molecule ligand to bind with E3 ligase and another ligand to recruit a target protein^16^. Consequently, the degradation of IDPs using PROTAC is still challenging as the lack of stable binding pockets presents a significant obstacle to develop high affinity small molecule ligands^17^. Although antibody degraders have been introduced in TPD, they are usually not cell permeable and, therefore, not ideal to degrade the large repertoire of IDPs located inside cells^18, 19^, or requiring invasive means like microinjection to access cytosolic antigens^11^.

A nanobody is a camelid-derived variable domain of heavy chain only antibodies (V_HH_). It is compact in size (∼15 kDa), binds with antigens with high specificity and strong affinity and can be produced at a low cost through bacterial expression^20^. Additionally, nanobodies are capable for binding with either structured or disordered proteins^21^. Due to the small size, nanobodies can be chemically or genetically engineered to be cell-permeable by attaching with appropriate cell-penetrating peptides^22, 23^. Recent studies showed that intrinsically disordered proteins are degraded by the endogenous 26S proteasome inside cells^24, 25^. Moreover, the disordered nature of IDPs may also favor the degradation through proteasomes which require ATP-dependent substrate unfolding before proteolysis^26^. Hence, we envisioned the possibility to design proteolysis targeting nanobody conjugate (PROTNC) for depleting IDPs. We will demonstrate the degradation of multiple IDP candidates of essential biological functions, like the end binding protein 1 (EB1)^27^, the antioncogenic protein p53^28^, and the microtubule nucleation factor TPX2^29^ (**Supplementary Table 1**).

## Results

### Design and generation of cell-permeable PROTNCs

First, we designed PROTNC which features a nanobody for recruiting a protein of interest, a AHPC ligand that targets the von-Hippel-Lindau (VHL) E3 ligase, and a cyclic cell-penetrating peptide (CPP) named cR_10_* through a disulfide bond (**Figure 1a**). We preferably employed a VHL ligand called AHPC because VHL was evaluated to exert a higher substrate specificity^30^. The CPP contains a cyclic-(KrRrRrRrRrRE)-NH_2_ (r: D-Arg; R: L-Arg) peptide with an N-terminal L-cysteine prepared through solid phase peptide synthesis (**Figure 1b**). This PROTNC is designed to cross the plasma membrane and recruit an IDP to VHL, leading to the ubiquitination of the IDP and its subsequent degradation by the proteasome (**Figure 1c**). As a proof of concept, we created a PROTNC called cR_10_*-SS-GBP-AHPC (cRGA), which carries a green fluorescent protein (GFP) binding protein (GBP), a nanobody that binds GFP with high affinity (K_d_ = 1.2 nM)^31^. Here, the use of a fluorescent protein (FP) as the substrate facilitates not only immunoblotting detection, but also fluorescence visualization. cRGA was assembled in two steps through coupling between Cys-AHPC ligand (**Supplementary Figure 1a**) with GBP-Intein (**I**) via expressed protein ligation (EPL) followed by disulfidization (**Supplementary Figure 1b-e**). We used both reducing and non-reducing SDS-PAGE to reveal that cR_10_* has been attached to GBP-AHPC (**II**) conjugate via a reductively cleavable disulfide bond (**Supplementary Figure 1e**).

**Figure 1.**
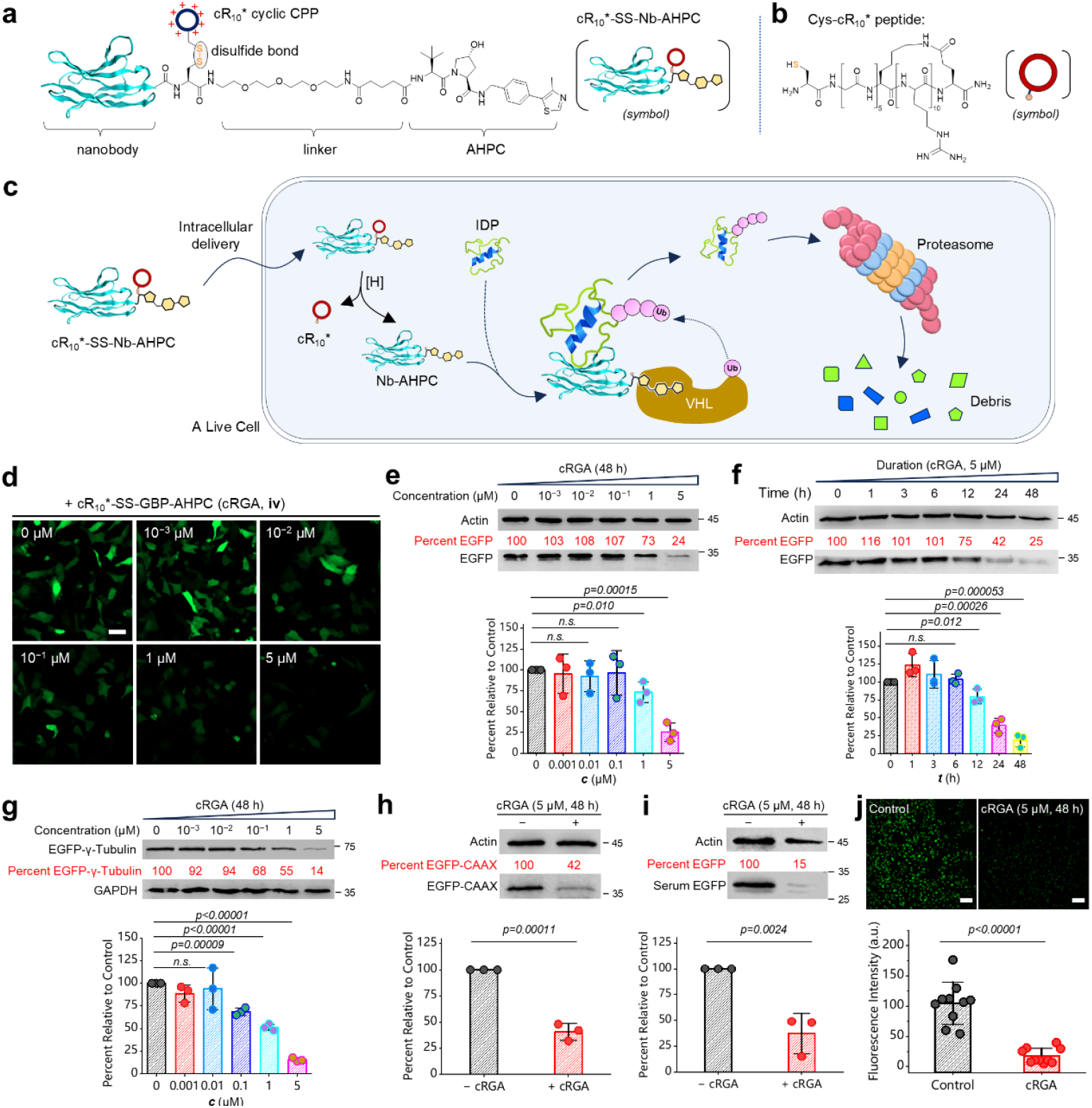
Design and characterization of PROTNC. **a**, Schematic view of the structure of a general PROTNC degrader which contains a nanobody module for binding with an IDP, a linker which contains biocompatible water-soluble polyethylene glycol (PEG) element, and a AHPC ligand for targeting to VHL E3 ligase; a symbolic view of a PROTNC is given at the right. **b**, Chemical structure of the cyclic cell-penetrating peptide CG_5_-cyclic-(KrRrRrRrRrRE)-NH_2_ (Cys-cR_10*_) (r: D-Arg; R: L-Arg); a symbolic view of it is given at the right. **c**, Schematic view of the mechanism of degradation mediated by PROTNC. **d**, Representative confocal micrographs of live HeLa cells expressing EGFP treated with gradient concentrations of cRGA; scale bar: 50 μm. **e**.WB analysis of the degradation of EGFP using gradient concentrations of cRGA. **f**, WB analysis of the degradation of EGFP in a time-dependent manner after adding cRGA (5 μM). **g**, WB analysis of the degradation of EGFP-γ-Tubulin using gradient concentrations of cRGA. **h**. WB analysis of the degradation of inner membrane protein EGFP-CAAX with or without adding cRGA (5 μM). **i**, WB analysis of the degradation of serum containing EGFP (5 μM) with or without adding cRGA (5 μM). **j**, Representative confocal micrographs of a large field of view with or without adding cRGA (5 μM); scale bar: 100 μm. Statistical quantifications were performed for results in **e-j**: n=3 experiments (**e-i**); n=10 fields (**j**); Student’s *t*-test was used (*n*.*s*.: non-significant).

### Cell-permeable PROTNC degrades intracellular, membrane-anchored, and serum EGFP and its fusions

Next, we evaluated that whether cRGA is able to degrade EGFP and its fused chimeras. To study this, we first showed that cRGA (**IV**) is cell-permeable via visualization of intracellular translocation of a fluorescently labeled cRGA (**Supplementary Figure 2**). Then, live HeLa cells expressing EGFP was treated with gradient dosages of cRGA for 48 h. Confocal micrographs revealed that a dose-dependent reduction of the fluorescence intensity of EGFP, particularly when the dosage of cRGA reaches 1 μM and above (**Figure 1d**). The degradation of the protein was further confirmed through Western blot (WB) analysis (**Figure 1e**). The rate of degradation was also examined, revealing that apparent degradation occurred after 12 h, with stronger depletion after 24 h (**Figure 1f**). Additionally, we observed that EGFP-fused protein, EGFP-γ-tubulin, can still be degraded by the same cRGA (**Figure 1g**). Furthermore, we conducted tests on an EGFP-fusion protein localized at the plasma membrane, and the WB result confirmed the successful degradation (**Figure 1h**). Finally, we were interested to see whether cRGA is able to degrade serum proteins because a PROTNC can complex with a protein and then be able to transport the substrate for intracellular degradation. For this, we supplemented cell culture medium with 5 μM of EGFP mixed with 1 equiv. of cRGA. Delightfully, both WB analysis and microscopic results confirmed the degradation of the serum EGFP (**Figure 1i-j**). Therefore, PROTNC is a robust degradation system that allows targeted degradation of intracellular, membrane-bound and extracellular proteins.

**Figure 2.**
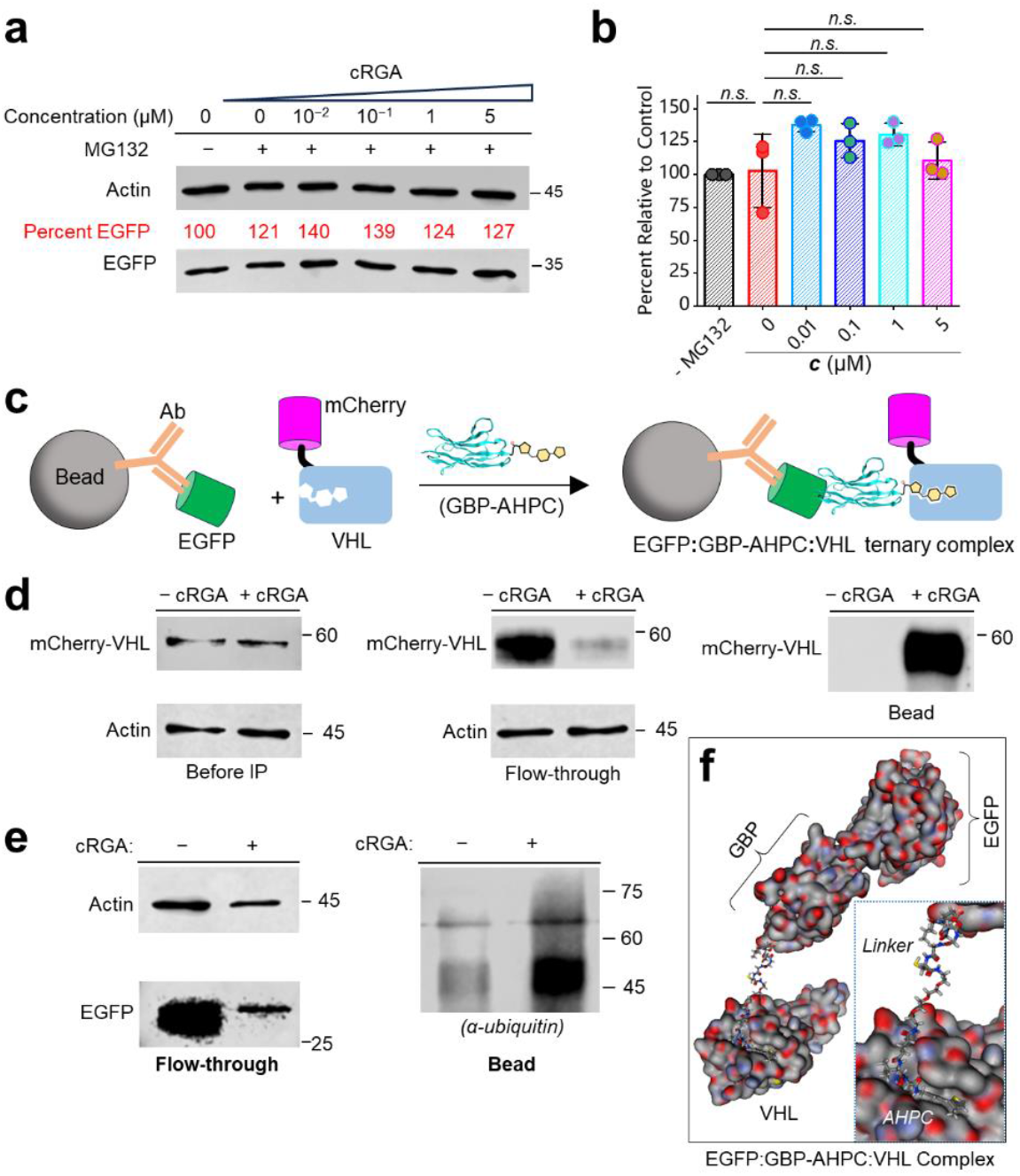
Mechanistic investigation reveals that PROTNC works through UPS degradation pathway. **a**, WB analysis revealed that EGFP in live HeLa cells was not degraded in the presence of MG132 (5 μM) after adding gradient concentrations of cRGA for 48 h. **b**, Statistic quantification of the WB results (n=3 experiments). **c**, Schematic view of the pulldown assay for demonstrating the formation of EGFP:GBP-AHPC:VHL ternary complex in the presence of GBP-AHPC; live HeLa cells coexpressing mCherry-VHL and EGFP were treated with cRGA (5 μM) and MG132 (5 μM) for 12 h, and then the cells were washed, lysed, and subjected to pulldown using GFP antibody coated manganic beads. **d**, WB of pulldown results: Left, the cell lysate input before incubation with α-GFP coated beads; middle, the follow-through after incubation with α-GFP coated beads; right, WB of the beads sample. **e**, HeLa cells co-expressing EGFP and Flag-ubiquitin were incubated with cRGA (5 µM) and MG132 (5 µM) for 12h; subsequently ubiquitinated proteins were immunoprecipitated by Flag-tag mAb coated beads and further analyzed by WB. **f**, Molecular operation environment (MOE) modeled 3D-structure of the EGFP:GBP-AHPC:VHL ternary complex; see Methods for more details.

### Design of a carrier-assisted PROTNC degradation platform

Although a PROTNC carries a cyclic CPP to enable intracellular delivery, the attachment of it through a disulfide bond complicates the overall preparation, and requires more purification steps. Inspired by the result of degrading serum-containing proteins (**Figure 1i-j**), we envisioned a carrier-assisted delivery platform to skip the attachment of cR_10_* (**Supplementary Figure 3**). We introduced an additional mCherry tag to a nanobody-AHPC conjugate and this is named as the “degrader” module which is not cell-permeable. The mCherry tag acts as both a fluorescent reporter allowing visualization of intracellular delivery, and as a binding tag for loading onto a carrier (**Supplementary Figure 3a**). Meanwhile, the cR_10_*-SS-RBP “carrier” module was generated which contains a mCherry red fluorescent protein binding protein (RBP) bridged with cR_10_* through a disulfide bond (**Supplementary Figure 3b**). Mixing of degrader and carrier in a one-to-one stoichiometry will give a complex that is able to enter live cells easily (**Supplementary Figure 3c**).

**Figure 3.**
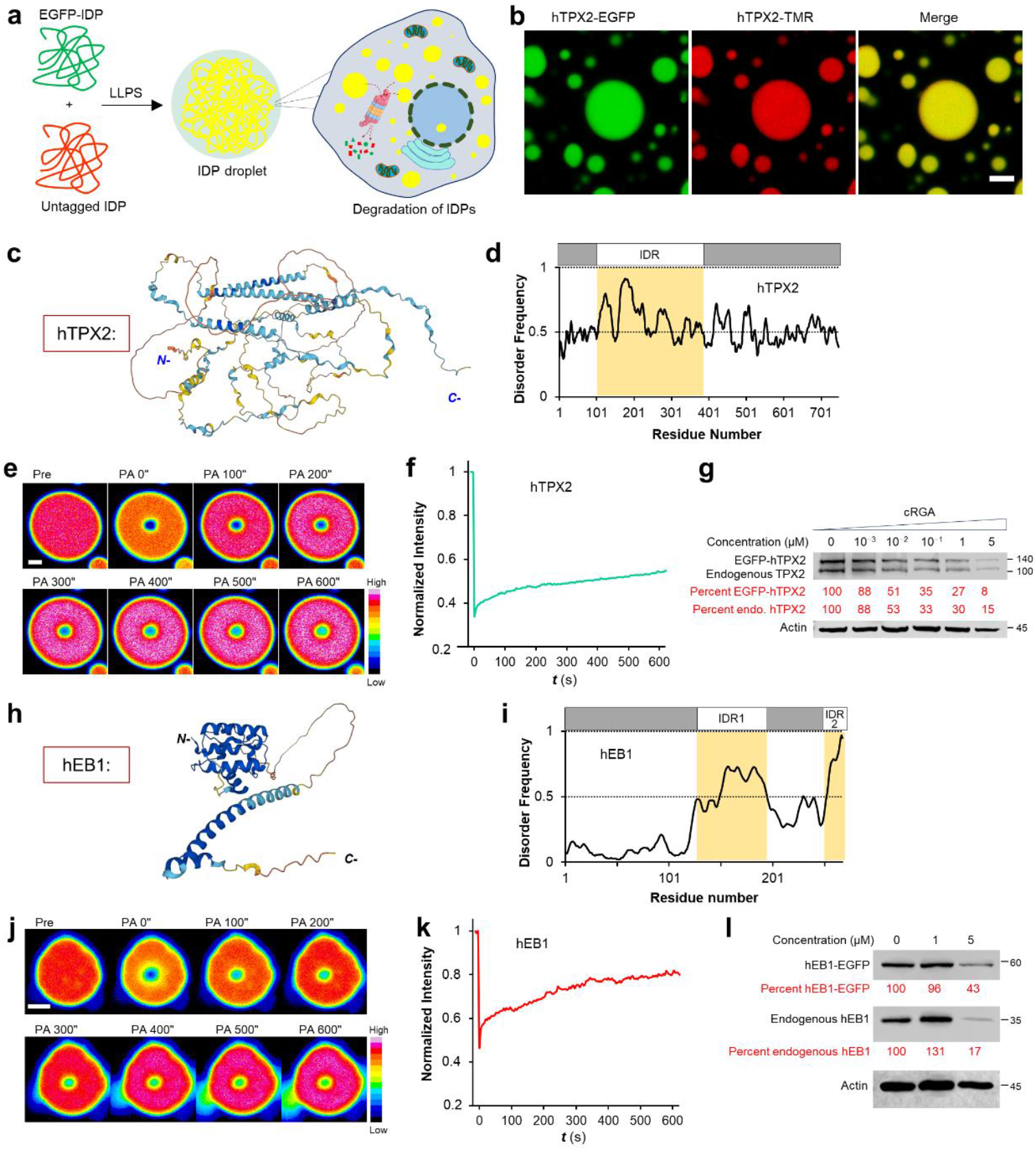
PROTNC degrades endogenous IDPs through co-degradation of FP-tagged IDPs based on the mechanism of co-condensate formation. **a**, Schematic view shows that a FP-tagged IDP together with an untagged IDP could form a co-condensate and got degraded together by PROTNC. **b**, Representative confocal micrographs revealed that hTPX2-EGFP forms con-condensates with TMR-labeled hTPX2; scale bar: 5 μm. **c**, AlphaFold2 predicted 3D structure of the disordered protein hTPX2. **d**, Secondary structure prediction using IUPred3 revealed that hTPX2 carries a large disordered region at the N-terminus. **e**, Representative confocal micrographs of the FRAP experiment of hTPX2-EGFP condensates; scale bar: 1 μm. **f**, Quantification of the fluorescence intensity at the region of photobleaching before and after photobleaching. **g**, WB analysis reveals that endogenous hTPX2 was degraded simultaneously with the depletion of EGFP-hTPX2 in live HeLa cells by cRGA for 48 h. **h**, ALphaFold2 simulated 3D structure of hEB1. **i**, Secondary structure prediction using IUPred3 revealed that hEB1 carries a large disordered region at the C-terminus. **j**, Representative confocal micrographs of the FRAP experiment of hEB1-mScarlet condensates; scale bar: 1 μm. **k**, Quantification of the fluorescence intensity of the region of photobleaching before and after photobleaching. **l**, WB analysis reveals that endogenous hEB1 was degraded along with co-depletion of hEB1-mScarlet in live HeLa cells after adding cRGA (5 μM) for 48 h.

To test our hypothesis, we evaluated the delivery system for the degradation of endogenous RhoA, a small GTPase that regulates cell morphology, motility, and transformation, using a nanobody against RhoA (i.e. RhoNb)^32^. Structural biology study^33^ and secondary structure prediction reveal that RhoA contains IDRs mainly at its C-terminus (**Supplementary Figure 3d**). For this, the degrader mCherry-RhoNb-AHPC (**VI**) (**Supplementary Figure 3e**) was generated via EPL between mCherry-RhoNb-Intein (**V**) and Cys-AHPC (**1**) (**Supplementary Figure 4**). Meanwhile, the carrier cR_10_*-SS-RBP (**IX**) was prepared via EPL between RBP-Intein (**VII**) and L-cysteine to produce RBP-Cys (**VIII**), followed by disulfidization coupling with Cys-cR_10_* (**Supplementary Figure 5**). We confirmed that the degrader is delivered into live cells in the presence of carrier using confocal microscopic imaging (**Supplementary Figure 3f-g**) and WB analysis (**Supplementary Figure 3h**). Afterwards, we demonstrated that this delivery PROTNC platform enables degradation of endogenous RhoA in a dose-dependent fashion (**Supplementary Figure 3i**). Hence, a carrier-assisted delivery PROTNC was introduced that facilitates the preparation of a PROTNC by waving the attachment of cR_10_* module.

**Figure 4.**
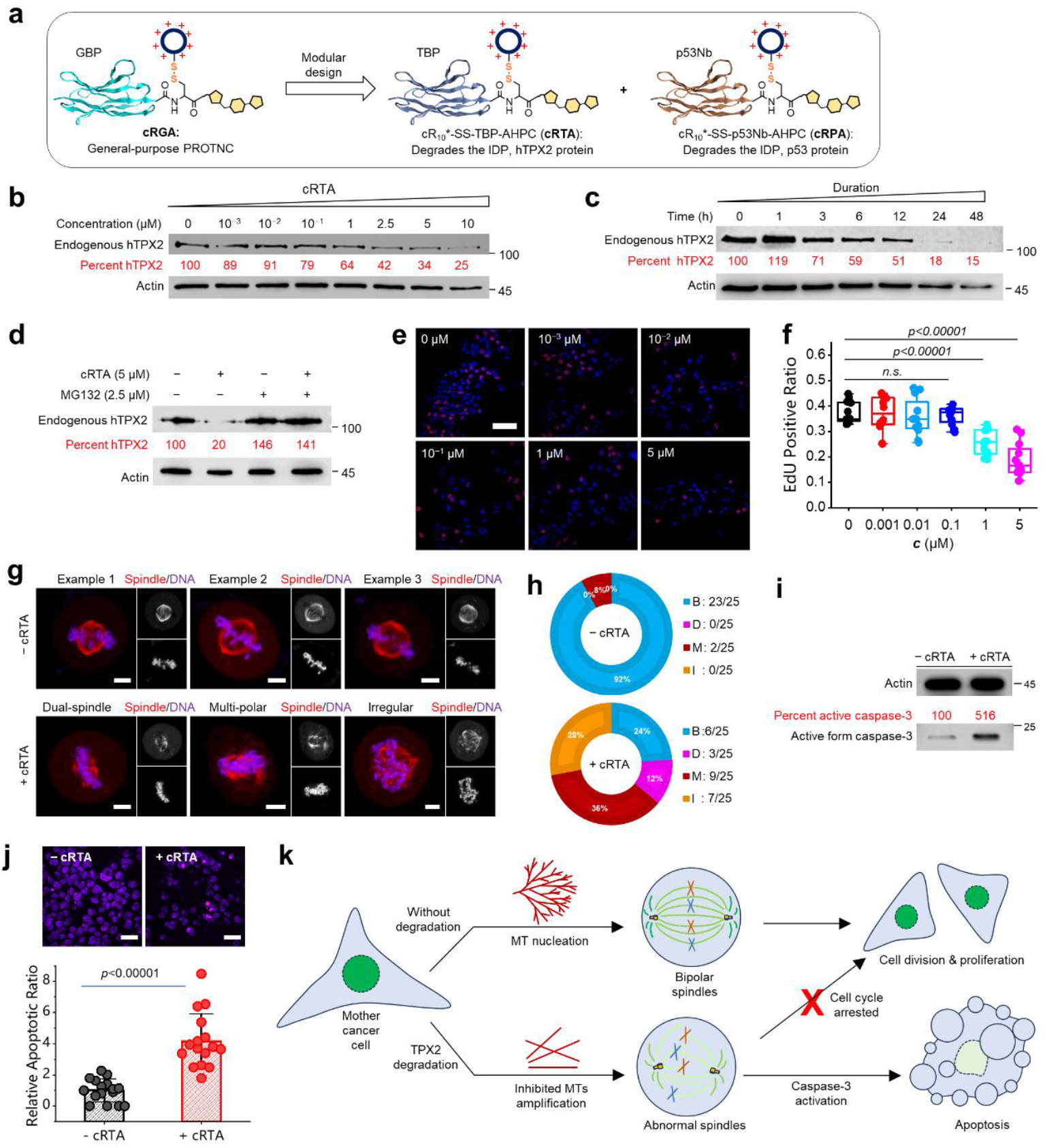
Design of PROTNCs for straightforward degradation of endogenous IDPs.**a**, Modular design of new PRONCs for degradation of IDPs like hTPX2 and p53. **b**, WB analysis of live HeLa cells treated with increasing concentrations of cRTA for 48 h revealed the clearance of hTPX2 in a dose-dependent fashion. **c**, WB analysis of live HeLa cells treated with cRTA (5 μM) for different durations revealed time-dependent degradation. **d**, WB analysis revealed that degradation of hTPX2 by cRTA (5 μM, 24 h) was inhibited in the presence of the proteasome inhibitor MG132 (2.5 μM, 24 h). **e**, Representative confocal micrographs of EdU proliferation assay of live HeLa cells treated with increasing concentrations of cRTA for 24 h; scale bar: 50 μm. **f**, Statistical quantification of the EdU positive ratio (n=12 fields); Student’s t-test was used. **g**, Representative micrographs of mitotic spindles captured in M-phase of live HeLa cells with or without treatment by cRTA (5 μM, 24 h); scale bar: 5 μm. **h**, Quantification of the percent of bipolar and non-bipolar/ multi-polar shaped spindles. **i**, Live HeLa cells treated with cRTA (5 μM, 24 h) was directed to cell apoptosis as evidenced by the enhancement of the key apoptotic factor, the active form of caspase-3. **j.**cRTA treated HeLa cells revealed the apoptotic phenotype of nucleus condensation as detected by Hoechst staining (up, scale bar: 50 μm); statistic quantification of the relative population of abnormal nuclei (down, n=16 fields); scale bar: 50 μm. **k**. A mechanistic scheme that explains how the inhibition of cancer cell proliferation occurs through the degradation of TPX2.

### Mechanistic investigation of the working principle of PROTNC

Then, we asked if the degradation proceeds through UPS pathway. In this regard, we added a classic proteasome inhibitor, MG132, into the cell culture medium and treated the cells with gradient concentrations of cRGA. WB analysis and statistical quantification revealed that the degradation of EGFP was fully inhibited (**Figure 2a-b**). Afterwards, we asked whether GBP-AHPC has directed EGFP to the E3 ligase VHL during the course of degradation. Therefore, we performed a pulldown assay using anti-GFP antibody (α-GFP) coated magnetic beads. Briefly, live HeLa cells coexpressing mCherry-VHL and EGFP were treated with cRGA (5 µM) and MG132 (5 µM) for 12 h, and then cells were washed, subjected to lysis, and the cell lysate was incubated with α-GFP coated beads (**Figure 2c**). WB revealed that the flowthrough has generally lost mCherry-VHL while the magnetic beads have recruited the mCherry-VHL (**Figure 2d**). Hence, we predicted that the addition of cRGA targeted EGFP to VHL E3 ligase inside living cells. Subsequently, we confirmed the ubiquitination of EGFP by probing the immunoprecipitated protein with an anti-ubiquitin antibody (α-ubiquitin), as shown in **Figure 2e**. Hence, PROTNC operates through the UPS pathway, involving the participation of VHL ligase and proteasomal degradation of the ubiquitinated protein substrate. Based on these results and the reported structure of an AHPC analogue bonded VHL^34^, we generated a 3D molecular modeling image of the EGFP:GBP-AHPC:VHL ternary complex that is formed during degradation using molecular operation environment (MOE) (**Figure 2f**).

### PROTNC degrades endogenous IDPs through co-degradation of tagged IDPs

After establishment of PROTNC platform, we moved forward to degrade IDPs. Most IDPs contain intrinsically disordered regions (IDRs) that render an IDP prone to undergo liquid-liquid phase separation (LLPS) forming so called lipid condensates or droplets^35^. Therefore, we envisioned the possibility to design a general pipeline for the degradation of endogenous IDPs via co-degradation of tagged IDPs (**Figure 3a**). We first looked at targeting protein for Xklp2 (TPX2), a recently identified intrinsically disordered protein with phase separation behavior^29^. Delightfully, we found that fluorescent protein (FP)-fused human TPX2 (hTPX2) indeed forms co-condensates with non-FP tagged hTPX2 (**Figure 3b**). Meanwhile, we performed 3D-structure simulation by AlphaFold2 (**Figure 3c**) and secondary structure prediction by IUPred3 (**Figure 3d**) to further confirm that hTPX2 is an IDP with disordered regions largely positioned at the N-terminal segment. Fluorescence recovery after photobleaching (FRAP) further demonstrated the phase separation behavior of hTPX2 (**Figure 3e-f**).

Therefore, it is possible that the degradation of FP-fused hTPX2 will also lead to the degradation of endogenous hTPX2. We therefore performed WB analysis and delightfully, both endogenous TPX2 and EGFP-TPX2 were degraded by cRGA even at a concentration below 1 μM (**Figure 3g**). To further validate this co-degradation strategy, we looked at another IDP called end binding protein 1 (EB1) which was recently reported to be an intrinsically disordered phase separation protein^27^. According to AlphaFold2 simulation and secondary structure prediction, we found that the C-terminal region of human EB1 (hEB1) is indeed largely intrinsically disordered (**Figure 3h-i**), and this protein shows phase separating behavior as evidenced by FRAP (**Figure 3j-k**). Finally, WB analysis demonstrated the degradation of endogenous hEB1 along with FP-tagged hEB1-mScarlet (**Figure 3l**). Therefore, we showcased a general strategy to degrade IDPs based on a mechanism of co-condensate formation.

### Degradation of an endogenous IDP by PROTNC that bears the respective nanobody

After demonstration of the general strategy, we were motivated to introduce PROTNCs capable of degrading an endogenous IDP in a straightforward fashion. This strategy will have the benefit to skip gene transfection and be invaluable in drug development. Here, we show that the GBP nanobody can be modularly exchanged to another nanobody, such as p53 nanobody (p53Nb) or TPX2 binding protein (TPB) nanobody, that directly binds with a respective endogenous IDP (**Figure 4a**). We first designed the PROTNC for degradation of p53, another intrinsically disordered protein^36^ with disordered regions located at both the N- and C-terminus (**Supplementary Figure 6a-b**). We designed the PROTNC called cR_10_*-SS-p53Nb-AHPC (cRPA), which carries a p53 nanobody (p53Nb), for targeted degradation of endogenous p53 (**Figure 4a, Supplementary Figure 6c**). Similarly, cRPA was prepared in two steps using the same modular preparation pipeline (**Supplementary Figure 7**). As a tumor suppressor, p53 keeps cells from growing or dividing too fast or in an uncontrolled way; one of the main pathways is to induce cancer cell apoptosis through activation of caspase-3^37^. The hemostasis of p53 was regulated by the endogenous Mdm2 E3 ligase which ubiquitinates it for downstream proteasomal degradation^38^ (**Supplementary Figure 6d**, up). Therefore, we were interested to see whether cRPA-based PROTNC degrades this anti-oncogenic target and triggers related cellular responses (**Supplementary Figure 6d**, down). According to WB analysis, live HeLa cells treated with cRPA for 24 h induces p53 degradation in a dose-dependent fashion (**Supplementary Figure 6e**). We also found that this degradation is related to cell apoptosis because the down-regulation of the key apoptotic marker caspase-3 was observed (**Supplementary Figure 6f**).

### Degradation of the oncogenic hTPX2 by PROTNC inhibits cancer cell proliferation

After demonstrating this straightforward strategy, we were primarily interested in the degradation of TPX2, a key microtubule nucleation factor that functions like a molecular glue to concentrate nucleation factors at the site of MT branching^29^. More importantly, the oncogenic TPX2 is of great therapeutic relevance because it is overexpressed in many cancer cells^39^. Therefore, we generated cR_10_*-SS-TBP-AHPC, i.e. cRTA, using the established procedure (**Supplementary Figure 8**). WB analysis revealed dose-dependent degradation of hTPX2 after adding cRTA for 24 h (**Figure 4b**). We found that the degradation of hTPX2 by cRTA (5 μM) becomes more complete along with longer incubation time (**Figure 4c**). We also validated that the degradation is UPS-dependent because the presence of MG132 inhibited the degradation (**Figure 4d**).

Subsequently, we investigated the inhibitory effect of degrading hTPX2 on cancer cell proliferation. The 5-ethynyl-2-deoxyuridine (EdU) assay revealed that HeLa cell proliferation was inhibited (**Figure 4e**), and this inhibition was more pronounced at higher concentrations of the drug (**Figure 4f**). Since TPX2 has been recognized as an essential factor in spindle assembly (**Supplementary Figure 9a**), we showed that depleting TPX2 abolishes microtubule nucleation which disrupts bipolar spindle assembly in Xenopus laevis egg extract (**Supplementary Figure 9**). We also found that the addition of cRTA caused the formation of a large portion of abnormal spindles including multi-polar, and remaining non-bipolar and irregular ones, whereas most spindles were bipolar without adding cRTA (**Figure 4g-h**). The proper assembly of mitotic spindle is essential for mitotic cell progression; failure of this process may direct cells to the fate of apoptosis^40^. We hence proceeded to determine apoptotic induction. It was observed that cells treated with cRTA showed a significant increase in active caspase-3 level, a key apoptotic maker (**Figure 4i-j**). Based on above results, we sketched a mechanistic scheme that explains how the inhibition of cancer cell proliferation occurs through the degradation of TPX2. This degradation disrupts proper bipolar spindle assembly by inhibiting microtubule nucleation, leading to the arrest of mitotic progression and ultimately inducing cell apoptosis (**Figure 4k**).

## Conclusion

In summary, we introduced proteolysis targeting nanobody conjugate (PROTNC) as a general and robust strategy for depleting intrinsically disordered proteins. PROTNC utilizes a nanobody for recruiting IDPs, a ligand such as AHPC to target an E3 ligase, and a cell-penetrating peptide for intracellular delivery. Firstly, we exploited the propensity of IDPs to form co-condensates through liquid-liquid phase separation, resulting in the degradation of endogenous IDPs along with the depletion of a FP-tagged IDP using a general-purpose degrader. Secondly, we introduced a straightforward degradation pipeline using a PROTNC that carries a nanobody specifically recognizes that particular IDP. We demonstrated the degradation of several disordered proteins, including hEB1, p53, and hTPX2. Hence, PROTNC is a valuable addition to the existing TPD toolbox, particularly suitable for degrading IDPs.

Importantly, we designed a PROTNC to degrade the oncogenic protein TPX2, an essential microtubule nucleation factor responsible for spindle assembly and cell division^41^. TPX2 is overexpressed in many cancer cells, and is considered as a promising target for anticancer drug development^39^. Unfortunately, due to its disordered and undruggable nature, previous studies mainly relied on genetic perturbations, such as RNA interference (RNAi)^42^ or others, to reduce TPX2 expression. Hence the direct biological impact of TPX2 depletion at protein level is largely elusive. In this study, we provided evidences for the first time that degrading TPX2 efficiently suppresses cancer cell proliferation and triggers apoptosis. Through mechanistic studies, we found that depleting TPX2 disrupts proper bipolar spindle assembly, leading to a halt in mitotic progression.

As a hybrid form of degrader, PROTNC takes the advantages of both small molecule PROTACs and antibody degraders^18, 19^. On one hand, PROTNC is cell-permeable being capable to degrade intracellular, membrane and serum proteins. On the other hand, it leverages the benefits of nanobodies to recognize and deplete either structured or disordered proteins. One limitation of PROTNC is the need for organic synthesis and bioconjugation during preparation. Nevertheless, thanks to its modular feature, a range of PROTNCs can be generated simply by coupling the same chemically synthesized ligand with a panel of nanobody fusions. Additionally, the carrier-assisted PROTNC platform also simplifies the implementation. In this paper, we have shown that the ubiquitin-proteasome system is well-suited for the targeted degradation of IDPs. However, it is important to note that alternative pathways, such as autophagy, have also been recently shown to degrade condensates in metazoan cells^43^. Therefore, future research can explore the use of these pathways in designing next generation degraders for IDPs. Overall, we anticipate that the PROTNC platform will greatly contribute to targeted protein degradation, benefiting both basic research and drug development in the future^44^.

## Methods

### Animal welfare

For Xenopus laevis (African clawed frog) husbandry, animal housing, maintenance, and egg harvesting were all carried out under approval from the IACUC of HIT with the permit number IACUC-2020020. Briefly, frog husbandry for both female (2-3 years old) and male xenopus laevis were conducted using a Xenopus laevis husbandry system supplied by a domestic manufacturer (LingYunBoJi, Beijing, P.R. China). Water quality (deionized water, ID-H_2_O), pH (7.2), temperature (18 °C), and conductivity (1600 μS/cm) etc. parameters were all set as advised by the manufacture’s manual, and Xenopus laevis were fed with qualified frog pellets twice per week. Male frogs were used for the preparation of sperm nuclei in spindle assay while female ones were used for inducing egg laying. Induction of frog ovulation was performed by sequentially injecting the female frogs with appropriate amount of PMSG (pregnant mare serum gonadotropin) for priming and hCG (human chorionic gonadotropin) for boosting with referring to CSH protocols (*Shaidani et al*., 2021).

### Mammalian cell culture

HeLa (#CL-0101) was obtained from Procell Life Science & Technology Co., Ltd. (Wuhan, P.R. China) while HeLa cells stably expressing mCherry-α-tubulin (#CY016) was obtained from Inovogen Tech. Co., Ltd. (Chongqing, P.R. China). The cells lines were short tandem repeat (STR) identified and proven to be HIV-1, HBV, HCV, mycoplasma, and other microorganisms free before culturing. Other reagents such as full DMEM (Dulbecco’s modified Eagle’s medium) and PBS (phosphate buffered saline) were also confirmed to be mycoplasma free before usage. HeLa cell culture was maintained at 37°C under 5 % CO_2_ in high glucose (4.5 g·L^−1^) DMEM (HyClone, #SH30243.01) containing 4 mM L-glutamine and sodium pyruvate and supplemented with additional 10 % fetal bovine serum (FBS) (HyClone, #SV30087.03), 1 % non-essential amino acid (NEAA, 100×), and 1 % penicillin-streptomycin (100×). Trypsin-EDTA (HyClone, #SH30042.01) and PBS (HyClone, #SH30256.01) were used in subculturing. HeLa cells were subcultivated in a ratio of 1:5∼10. HeLa cells stably expressing mCherry-α-tubulin was maintained at 37°C under 5 % CO_2_ in minimum essential medium (MEM) (PriCella, #PM150414) containing 4 mM L-glutamine and sodium pyruvate and supplemented with additional 10 % fetal bovine serum (FBS) (HyClone, #SV30087.03), 1 % non-essential amino acid (NEAA, 100×), and 1 % penicillin-streptomycin (100×), 0.1 % puromycin (Shanghai MaokangBio, #MS0011-100MG). Trypsin-EDTA (HyClone, #SH30042.01) and PBS (HyClone, #SH30256.01) were used in subculturing. mCherry-α-tubulin HeLa cells were subcultivated in a ratio of 1:2∼3.

### Plasmid construction

Plasmid vectors, such as pTXB1, pET28a(+), EGFP-C1 and EGFP-N1 were obtained from commercial vendors. These parental vectors may be further engineered, such as introducing a His_6_-or His_8_-affinity tag, insertion of a TEV (ENLYFQ↓G) or TEV’ (ENLYFQ↓C) protease cleavage site, alternation of restriction cleavage sits, or replacing EGFP by other fluorescent proteins, e.g. mCherry, etc. to give modified versions of the parental vector for cloning. Subcloning, Gibson cloning, or modified Gibson cloning methods were employed to construct the desired plasmids. For subcloning, fragments of interest were directly cut from the parent plasmid using appropriate restriction enzymes, or amplified by PCR from plasmids containing the desired genes using hyPerFUsion high-fidelity polymerase (APExBIO, #1032,), gel purified, digested with restriction enzymes and purified again. The gene fragments were ligated into appropriate vectors using T4 DNA ligase. Multiple fragments were assembled by stepwise subcloning or one-step multi-fragment Gibson cloning. Genes of interest were obtained via custom gene synthesis from Comate Bioscience Co., Ltd. (Changchun, P.R. China) or Ruibiotech (Beijing, P.R.China). These genes include *E. coli*. codon optimized *homo sapiens* TPX2 (hTPX2), nanobody genes, and others. None-codon optimized TPX2 gene was amplified from the plasmid pLenti-EF1a-EGFP-P2A-Puro-CMV-TPX2-3Flag purchased from (Shanghai Obio Technology Co., Ltd., #H10559). Alternatively, plasmids containing the desired genes can be purchased from Miaoling Plasmid Sharing Platform if applicable.

### Transfection

Transient transfection was typically performed in an 8-well (#155409) Lab-Tek®II imaging chamber from Thermo Scientific using Lipo8000™ transfection reagent from Beyotime Biotechnology (#C0533). Typically, 0.25 μg DNA was dissolved in 12.5 µl gibco opti-MEM (Life technologies, #31985-062) and then 0.4 μl Lipo8000™ transfection reagent was added and mixed via gentle pipetting. Then this mixture was added into an imaging chamber well seeded with 2.5×10^4^ cells that were already adhesively attached on the bottom in 250 μl full DMEM. The cells were maintained under 5% CO_2_ at 37 °C for around 2 h. Then the medium was replaced by warm full DMEM and the cells were further incubated under 5% CO_2_ at 37 °C for over 20 h. For co-transfection of more than one plasmid, the quantity of DNA used in this protocol implies the total amount of plasmids.

### Confocal microscopy

Live cells were imaged in phenol red free Dulbecco’s Modified Eagle Medium (Life Technologies, #21063-29) supplemented with additional 10 % FBS, 1 % sodium pyruvate, 1 % NEAA, 1 % penicillin-streptomycin and 15 mM HEPES-Na at 37° under 5 % CO2. Microscopy was performed using Nikon A1 ECLIPSE T*i*2 inverted confocal microscope. The microscope was equipped with four lasers (405 nm, 488 nm, 561 nm, and 640 nm), allowing the acquisition of confocal fluorescence data for four different excitation wavelengths. For detection of blue (excited by 405 nm laser) or far red (excited by 640 nm) fluorescence signal, PMT (photomultiplier tube) detectors were used; while for the detection of green (excited by 488 nm) or red (excited by 561 nm) fluorescence signal, the more advanced GaAsP detector with even higher sensitivity will be used. Most images presented in this article were acquired with a 60× oil objective lens (APO 60×/1.40 oil) having a numerical aperture of 1.4. Alternatively, a 40× objective lens (Plan Apo 40×/0.95) having a numerical aperture of 0.95 can be used, for example when capturing images with larger field of views. In most cases, the basic imaging setup were configured with typical parameters set as follows: Scan speed 0.5, number of averaging 4, scan line mode in one-way scan direction. For fluorescence recovery after photobleaching (FRAP) associated microscopic imaging experiments, confocal microscopy was performed using Zeiss LSM 880 inverted confocal laser scanning microscope equipped with Zeiss Plan-APOCHROMAT 63×/1.4 oil DIC objective having a numerical aperture of 1.4.

### Co-condensate formation assay and FRAP experiment

Pure protein samples were first buffer exchanged to pH 7.4 buffer containing 50 mM Tris/HCl, 250mM NaCl, and 10% glycerol. And after high-speed centrifugation, the supernatant was collected and the protein concentration was determined using an ultra-micro ultraviolet-visible spectrophotometer via measurement of A280. In the co-condensate formation assay, fluorescent protein (FP)-fused intrinsically disordered protein (IDP) and the corresponding fluorophore-labeled IDP were mixed in a 1:1 stoichiometric ratio at room temperature to a final concentration of 10 μM. Afterwards, PEG8000 (Aladdin, #P103734-250g) was added to a final concentration of 10 %, and the reaction solution was incubated on ice for 2 minutes to allow co-condensate formation. Subsequently, the sample was loaded onto a clean glass slide, covered with a coverslip, and subjected to confocal microscopic imaging and fluorescence recovery after bleaching (FRAP) analysis.

Confocal imaging and FRAP analysis were conducted using a Zeiss LSM 880 confocal microscope equipped with a 63× oil objective (Zeiss Plan-APOCHROMAT 63×/1.4 oil DIC) with a numerical aperture of 1.4. For photobleaching of mScarlet, a 543 nm laser at 60% power (170 cycles) was used while for photobleaching of EGFP, a 405 nm laser diode was used at 90% power (20 cycles). Before photobleaching, several time-lapse images were acquired and after photobleaching, time-lapse images were recorded for a duration of 13 min with a 2-second interval.

### EdU cell proliferation assay

The EdU cytotoxicity/ cell proliferation assay was performed using an EdU cell proliferation detection kit (RiboBio, #R11053.9). Briefly, 1×10^4^ HeLa cells harvested at exponential phase were seeded in a 96-well plate with transparent glass bottom (Shanghai JingAn Biological, #J04961) suitable for high resolution confocal imaging. Afterwards, cells were allowed to grow overnight. Drug solutions at specified final concentrations in full DMEM was used to treat the cells. After a given incubation time, the drug solution was replaced by EdU solution at a final concentration of 50 μM. It was incubated for 2 hours at 37°C under 5% CO_2_ as recommended for general cancer cell lines. Then, each well was washed by PBS (2×5 min) to remove the excess of EdU, added with 100 μl of fixative solution (4% PMA in PBS) and incubated for 30 min at room temperature. Then 100 μl of 2 mg·ml^−1^ glycine solution was added to each well and shaken at RT for 5 min to quench the fixative. Glycine solution was removed and each well was washed by 200 μl PBS and shaken at RT for 5 min. PBS was removed and each well was added with 200 μl cell permeabilization solution (0.5% TritonX-100 in PBS) and shaken at RT for 10 min. The fixed cells were further washed by PBS (1×5 min) before labeling.

Before fluorescent labeling by click reaction, 1× Apollo labeling solution that contains the red color Apollo567 dye (RiboBio, #C10310-1), catalyst and other necessary reagents were freshly prepared according to the manufacture’s guidance. For example, 1 ml of 1× Apollo labeling solution could be prepared by sequentially adding 938 μl DI-H_2_O, 50 μl Apollo reaction buffer (reagent B), 10 μl Apollo catalyst solution (Cu^2+^, buffer C), 3 μl Apollo 567 dye (reagent D) and ∼9 mg Apollo additive (sodium ascorbate, reagent E). 200 μl freshly prepared 1× Apollo labeling solution was added into each well, shielding from light, and shaken at RT for 30 min to complete the click labeling. Labeling solution was removed, and the cells in each well were washed by permeabilization solution (0.5% TritonX-100 in PBS) again (3×10min). Permeabilization solution was removed and the cells were wash by PBS (1×5 min). Finally fresh PBS was added and the labeled cells were ready for confocal microscopy imaging. Hoechst could be used to label the nucleus if necessary.

### Immunodepletion (ID)

In a typical ID experiment, 32 μl of Protein A/G MagBeads (IP grade) (YEASEN, #36417ES03) were coupled with xenopus TPX2 (xTPX2) polyclonal antibody (∼ 23 μg) according to manufacturer’s protocol. xTPX2 polyclonal antibody was generated by immunization of Rabbit using the C-terminal fragment of xTPX2 protein, affinity purified via Protein A first, then using the antigen. The antibody-coated beads were suspended in 32 μl of CSF-XB buffer and spitted to three equal portions. Each bead portion was removed of CSF-XB buffer prior to egg extract addition. Then 30 μl of freshly prepared egg extract was mixed with one portion of bead followed by gentle pipetting to resuspend the beads. The suspension was incubated on ice for around 10-15 min, and then retrieved of beads using a magnetic particle concentrator (MPC) for 5-10 min. This ID process was repeated two additional times to completely deplete xTPX2 in the extract. Western blot using the same antibody for ID can be used to confirm the full depletion of xTPX2.

### Immunoprecipitation (IP)

HeLa cells were seeded in 24-well cell culture plates at a density of 8×10^4^ cells/well in cell culture medium overnight to allow cell adhesion. The next day morning, the cells were transfected with the indicated plasmids for 24 h. Afterwards, HeLa cells were treated with 5 µM MG132(Beyotime, #S1748-5mg) and 5µM cRGA for 12h. Then cells were washed once with PBS and lysed in radioimmunoprecipitation assay buffer (RIPA) buffer (Biosharp, #BL504A) supplemented with additional 1% PMSF (Solarbio, #P3840) on ice for 10 min. The cell lysate was then centrifuged at 17,320 g at 4 °C for 10 min and the supernatant was collected for subsequent immunoprecipitation. IP-grade Protein A/G MagBeads (Yeasen, #36417ES08) were first coated with EGFP mAb (ZENBIO, # R24437) or Flag-tag mAb (Beyotime, #AF519) according to the manufacturer’s protocol. The beads were then retrieved using a magnetic particle concentrator (MPC), gently mixed with the freshly prepared cell lysate supernatant, and slowly rotated at 4 °C overnight. The next day, the beads were retrieved using MPC, washed three times by 1× TBST, and boiled with 50 μl 2× SDS-PAGE sample loading buffer at 90 °C for 5 min. After removing the beads using the MPC, the supernatant was subjected to Western blot (WB) analysis. For WB, 10 μl of the sample was separated using SDS-PAGE, transferred to a nitrocellulose membrane, and probed with appropriate antibodies.

### Western blot (WB) analysis

HeLa cells were plated in 24-well plate at a density of 6×10^4^ cells per well, and then left to grow overnight to allow cell adhesion before drug treatment at the specified concentrations. For the degradation of overexpressed protein such as EGFP and EGFP-fused proteins, cells were first subjected to transfection using respective vectors before adding drugs. Unless otherwise specified, after 48 hours or specific time, cells were washed (2×PBS), collected using 2× SDS-PAGE loading buffer (120 μl/well for 24-well plates, 40 μl/well for 8-well chamber), and then boiled at 90 °C for 5 minutes before subsequent gel electrophoresis and WB analysis.

For WB analysis, cell lysate samples were first subjected to 10%, 12% or 15% SDS-PAGE gel electrophoresis (180 V, 45 min), transferred to nitrocellulose (NC) membrane (PALL, #66485) on ice applying constant 400 mA current for 30 min in rapid WB transferring buffer (Genefist, #GF1816). The NC membrane was blocked using 5 % skim milk (Biosharp, #BS102) in 1× TBST at room temperature for 1.5 h. Then, the blocked NC membrane was labeled by primary antibodies diluted in 5 % skim milk at 4 °C overnight. These primary antibodies include: GFP-tag rabbit mAb antibody (Zenbio, #R24437) diluted at a ratio of 1:1000, mCherry-Tag antibody (BIOSS, #bs-41161R) diluted at a ratio of 1:2000, RhoA rabbit pAb antibody (Zenbio, #346086) diluted at a ratio of 1:1000, p53 rabbit mAb (Zenbio, #R25247) diluted in a ratio of 1:1000, TPX2 rabbit mAb (Zenbio, #R27376) diluted in a ratio of 1:500, EB1 polyclonal (Proteintech, #17717-1-AP) diluted in a ratio of 1:500, rabbit anti-caspase-3 polyclonal antibody (Bioss, #bs-0081R), β-actin rabbit mAb antibody (ABclonal, #AC026) diluted at a ratio of 1:50000, and GAPDH polyclonal antibody (Bioworld, #AP0063) diluted at a ratio of 1:5000. The next day, NC membrane was washed by TBST (3×5min), incubated with secondary antibody using HRP-conjugated goat anti-rabbit IgG antibody (Zenbio, #511203) diluted in 5 % skim milk at a ratio of 1: 5000 at room temperature for 1h, and washed by TBST (3×10min) before imaging. For WB signal detection, the membrane was developed with a mixture of high-sensitivity luminescent liquid (Biosharp, #BL523B). Densitometry of developed bands was measured and analyzed using LI-COR Odyssey Fc imaging system

### Spindle assembly and microtubule nucleation assay in egg extract

Xenopus laevis egg extract was prepared from freshly ovulated frog eggs in an 18 °C room with referring to the generally adopted protocols (*Hannak & Head, Nat. Protoc*., 2006, 1, 2305). The freshly prepared egg extract was added with 10 mg·ml^−1^ LPC (leupeptin, <u>p</u>epstatin, <u>c</u>hymostatin, each final 20 μg·ml^−1^) protease inhibitors and 10 mg·ml^−1^ cytochalasin D (final 20 μg·ml^−1^) but no energy mix, gently mixed and put on ice prior to use. Immunodepletion, spindle assay or microtubule nucleation assay should be conducted as soon as the egg extract was prepared. Sperm nuclei was prepared from male frogs according to CSH protocols (*Hazel & Gatlin*, 2018). For spindle assay, 0.25 μl sperm nuclei, 0.33 μl of 2 mg·ml^−1^ HiLyte 647 porcine brain tubulin (Cytoskeleton, Inc., #TL670M-A/B), 0.25 μl of 5 mg·ml^−1^ EB1-mCherry and 0.25 μl of 100 μg·ml^−1^ DAPI were added into 8 μl egg extract. The egg extract was gently mixed and then loaded into a self-prepared glass slide featuring sample channels. The extract mixture was incubated at 18 °C and spindle structures can be visualized using a confocal laser scanning microscope after 30 min. For microtubule nucleation assay, 0.3 μl EB1/vanadate (1 mg·ml^−1^ EB1-mCherry & 10 mM sodium vanadate) and 0.33 μl HiLyte 647 porcine brain tubulin (2 mg·ml^−1^) were added into 8 μl extract; the extract mixture was gently mixed and then immediately loaded into a glass slide featuring sample channels. The extract mixture was incubated at 18 °C and microtubules will appear gradually, which can be visualized using a high-end confocal laser scanning microscope.

### Design and preparation of the cyclic cell-penetrating peptide Cys-cR10*

The cyclic Cys-cR10* peptide was designed with a cyclic rR ring (r = D-Arg, R = L-Arg) plus a (Gly)_5_ linker with a N-terminal free cysteine and a C-terminal -CONH_2_ group. The peptide was synthesized via standard solid phase peptide synthesis from Rink amide resin. After the synthesis of liner R_10_* fragment, intramolecular cyclization was performed to bridge the Lys side chain (-NH_2_ group) and Glu side chain (-COOH group). Afterwards, Cys-(Gly)_5_ tail was sequentially added to the cyclic-R_10_* moiety followed by TFA deprotection and HPLC purification. Cys-cR_10_* peptide was obtained in a purity of 95.1 % and confirmed using mass spectrometry. C_84_H_160_N_50_O_19_S, exact mass: 2205.28, M.W.: 2206.56; found *m/z* 736.4 [M+3H]^3+^, 553.1 [M+4H]^4+^, 442.3 [M+5H]^5+^, 368.8 [M+6H]^6+^.

### Protein expression and purification

#### General protocol

pTXB1 vector was used to express intein-tag fused nanobody chimeras for expressed protein ligation (EPL) while pET28a(+) or modified pET28a(+) vectors were used in expression of other protein chimeras. These plasmids for protein expression were first transformed into *E. coli* Rosetta 2a cells and the transformants were selected on ampicillin (100 mg·L^−1^) or kanamycin (50 mg·L^−1^) agar plates depending on the antibiotic resistance of the plasmids. A single colony was used to inoculate 50-100 ml of LB medium containing 100 mg·L^−1^ ampicillin or 50 mg·L^−1^ kanamycin and shaken at 240 rpm for 8-10 hours or overnight at 37 °C. 30-50 ml of the preculture was used to further inoculate ∼1.8 L fresh LB medium containing 100 mg·L^−1^ ampicillin or 50 mg·L^−1^ kanamycin, and additional chloramphenicol (33 mg·L^−1^). The absorbance at 600 nm (OD600) of the inoculated culture should be controlled between 0.05 to 0.1 in this inoculation step. Then the culture was shaken at 180 rpm at 37°C for a few hours (typically 2-3 h) until OD 600 reached 0.5-0.6. Then 0.5 ml isopropyl β-D-thiogalactoside (IPTG) stock solution (1 M) was added (final ∼0.27 mM) to induce protein expression at 37 °C for 5 h, or at 16 °C overnight. Sometimes protein expression time and temperature needed to be optimized in order to achieve an optimal expression for some particular proteins.

Later, cells were harvested by centrifugation at 13881× g, at 4 °C for 15 min and washed once with PBS (4149× g, 10 min). The bacterial pellet was resuspended in lysis buffer (pH 8.0, PBS supplemented with additional 0.5 M NaCl, 3 % glycerol, w/o 3 mM β-mercaptoethanol (BME), 1 mM phenylmethylsulfonyl fluoride (PMSF). For relatively smaller volumes of bacterial cell suspensions (< 40 ml), bacterial cells were typically lysed via ultra-sonification at 80 W for 30 min or 60 W for 45 min (1 s sonification followed by 3 s interval) on ice. For batch processing or larger volumes of cell suspensions, cells were typically lysed using ultra-high-pressure homogenizer cooled by a bench chiller for 2-3 cycles under 800-900 bar at 4 °C. The lysate was cleared by high-speed centrifugation (74766× g, 45 min, 4 °C) and the supernatant was loaded onto a gravity Ni-NTA column (2-5 ml resin). The Ni-NTA column was washed and then the His-tag fused protein was eluted using step-gradient of imidazole (50, 100, …, until 500 mM) solutions. Alternatively, GE ÄKTA Pure machine equipped with a HisTrap FF column was used to purify His-tagged protein via gradient elution (0 → 500 mM imidazole) by combining buffer A (pH 8.0 PBS, 0.5 M NaCl, 3 % glycerol, w/o 3 mM BME) and buffer B (pH 8.0 PBS, 0.5 M imidazole, 0.5 M NaCl, 3 % glycerol, w/o 3 mM BEM). Ionic exchange or size exclusion chromatography may be further applied if additional purifications are necessary. The obtained proteins were typically concentrated, buffer exchanged in buffer A, aliquoted, snap frozen in liquid nitrogen, and stored under -80 °C.

### General EPL protocol for the preparation of nanobody-AHPC or nanobody-Cys conjugate

Proteins to be ligated were expressed as a fusion chimera with a C-terminal Mxe GyrA intein tag by cloning the respective gene into pTXB1 vector. Afterwards, this fusion chimera was expressed, purified and buffer exchanged in Buffer A (pH 8.0 PBS, 0.5 M NaCl, 3 % glycerol). Typically, the Mxe GyrA intein fusion protein was reconstituted to around 15 mg·ml^−1^ before ligation. To initialize the ligation reaction, 1/2 volume of pH 8.0 sodium 2-mercaptoethanesulfonate (MENSNa) stock solution (2 M) was added as the intein cleavage reagent and 1/2 volume of pH 8.0 4-mercaptophenylacetic acid (MPAA) stock solution (1.1 M) was added as the catalyst. Finally, Cys-AHPC (15 mM/ DMSO) was added to the reaction solution at a final concentration of 0.75 mM. The reaction mixture was incubated at 4 °C for 3 days. The ligated product was then purified via step-gradient (0 → 500 mM imidazole) reverse gravity Ni-IMAC chromatography using high affinity Ni-charged Resin FF (GenScript, #L00666-25,). In this process, cleaved intein-His^6^ fragment will bind onto the resin and pure nanobody conjugate will be eluted out. This process will typically generate nanobody-AHPC fusions with sufficient purity for the next step. Otherwise, further size exclusion chromatography purification can be used to further purify the conjugate.

### Stepwise protocol for assembly of PROTNCs

➢ EPL: 0.9 ml of nanobody-intein-His_6_ chimera at around 15 mg·ml^−1^ concentration in a 2 ml Eppendorf tube was added with 0.45 ml MENSNa (pH 8.0, 2 M) and 0.45 ml MPAA (pH 8.0, 1.1 M) stock solutions. Afterwards, 90 μl of Cys-AHPC (15 mM/DMSO) stock solution was added at final concentration around 0.75 mM. The reaction solution was degassed via sonification, charged with argon, sealed, shielded from light using aluminum foil, and incubated at 4 °C for 3 days to accomplish EPL.
➢ Reverse Ni-IMAC purification: The reaction mixture was subjected to Ni-NTA column purification to separate nanobody-AHPC conjugate. Pure conjugate fractions were combined, buffer exchanged in BME-free elution buffer (pH 8.0 PBS, supplemented with additional 3%v glycerol and 0.5M NaCl), degassed via sonication, added with 2 equiv. TCEP (20 mM stock), charged with argon, incubated at 4 °C for 45 min. Subsequently, 10 equiv. of DTNB (100 mM) stock solution in 0.5 M Na_2_HPO_4_ buffer was added portion-wise (2 equiv. × 5). The reaction solution will immediately turn yellowish, and the reaction solution was incubated at 4 °C under argon for 60 min. **Abbreviation:** IMAC, immobilized metal affinity chromatography.
➢ Disulfidization coupling: Buffer exchange into pH 9.0 disulfidization buffer (50 mM HEPES, 0.5 M NaCl) for three times, degas via sonification, add around 3 equiv. of Cys-cR_10_* (15 mM/ DMSO stock), degas via sonification, charged with argon, and incubate on ice overnight to finish the coupling. The next day, the reaction solution was buffer exchanged two times into PBS, analyzed via non-reducing SDS-PAGE, aliquoted, snap frozen in liquid nitrogen, and stored under -80 °C before use.

### Molecular modeling of the EGFP:GBP-AHPC:VHL ternary complex

Molecular modeling of the 3D-structure of this ternary complex involves three main steps. **Step I:** Modeling of the AHPC:VHL binary complex. Since the crystal structure of AHPC bond to VHL is currently unavailable, it is necessary to first generate a simulated 3D-structure of the bond form. To do this, molecular modeling was performed using the Molecular Operation Environment (MOE) software. Briefly, the pdb file (3ZRC) of the VHL E3 ligase complex with a highly potent inhibitor (an AHPC analogue) was imported into MOE (file/open). Then the inhibitor was modified bond-by-bond to AHPC using the molecular builder function (system/builder). Subsequently, the complex underwent hydrogen addition (compute/protonate 3D), partial charge calculation (compute/partial charges), and energy minimalization (compute/energy minimized) to produce the simulated 3D-structure of the AHPC:VHL binary complex. This was saved as a .moe file. **Step II:** Modeling of the EGFP:GBP-linker binary complex. The pdb file (3OGO) of the GFP:GBP binary complex was imported into MOE (file/open). Then the mutations (F64L/S65T) were generated using the molecular builder (system/builder) to convert GFP to EGFP while the PEG linker was attached to GBP to produce GBP-linker. The resulting complex was subjected to hydrogen addition, partial charge calculation and energy minimization as described in the preivous step to generate the modeled 3D-structure of the EGFP:GBP-PEG-linker binary complex. This was saved as a .moe file. **Step III:** Generation of EGFP:GBP-AHPC:VHL 3-D structure. In this final step, the two binary complex files generated in the previous steps are imported into MOE. Appropriate rotation and x/y/z movement are performed to position the AHPC:VHL binary complex and the EGFP:GBP-PEG-linker binary complex close to each other. The linker end should be positioned near the VHL ligand, maintaining a distance of a single covalent bond in order to facilitate the connection. The linker end and AHPC ligand are connected using the molecular builder (system/builder), and the resulting ternary complex is then further energy minimized to obtain the final simulated 3D structure of the EGFP:GBP-AHPC:VHL ternary complex. The VHL-linker part is depicted in sticks (view/model/stick) and colored in element (view/color/element) while the protein domains are represented in “Gaussian Contact” surface and colored as “atom color” (compute/surface and maps). The file is saved as a .moe for potential further operations and as .png for image visualization.

### 3D-Structure simulation of IDPs and prediction of IDRs

The crystal structures of full-length intrinsically disordered proteins (IDPs) are often unavailable. Therefore, in this study, the 3D structures of IDPs were simulated using AlphaFold2, which is accessed through the UniProt online database (www.uniprot.org). For predicting intrinsically disordered regions (IDRs), various methods are currently available. This study primarily relies on the results predicted by IUPred3 (https://iupred.elte.hu) online, as they are consistent with the predictions made by AlphaFold2. An alternative method, PONDR (Predictor of Natural Disordered Regions: www.pondr.com), can also be used for predicting IDRs, but its results typically cover a larger sequence. Hence, in this study, the regions predicted as IDRs by PONDR but not by IUPred3 are referred to as semi-disordered regions.

### Image analysis

Microscopic images were analyzed and processed with ImageJ/Fiji and prepared for presentation using Microsoft Office PowerPoint. Image manipulations were restricted to adjustment of brightness level (i.e. linear stretch), background subtraction, cropping, rotating, scaling, and false color-coding using Look-Up Tables (LUT).

### Statistics and reproducibility

All microscopic imaging experiments were representative of at least three independent repeats if not otherwise stated; representative SDS-PAGE images were from at least three independent repeats with similar results; representative confocal microscopic images were from at least ten independent cells with similar results; IP experiment was performed once which already provided clear results. No randomization nor blinding was used in this study. Origin and Microsoft Excel were used for plotting, data fitting, graphing and statistical analysis. All box plots show mean (square), median (bisecting line), bounds of box (75^th^ to 25^th^ percentiles), outlier range with 1.5 coefficient (whiskers), and minimum and maximum data points (lower/ upper whiskers). Student’s *t*-tests were used to compare two experimental conditions. Unless otherwise specified, one-sided unpaired *t*-tests were performed, as for example cells with or without drug treatment. When necessary, stars were used to denote *P*-values for indicated statistical tests (*: *P*<0.05; **: *P*<0.01; ***: *P*<0.001; ****: *P*<0.0001). Exact *P*-values were indicated for critical experiments.

## Acknowledgement

We thank the Core Facilities at HCLS and School of Life Science and Technology in Harbin Institute of Technology. This work was supported by the funding from Harbin Institute of Technology, Overseas Outstanding Young Talents Program of China, National Natural Science Foundation of China (grant No. 32071410 to X.C.), and Natural Science Foundation of Heilongjiang Province of China (grant No. YQ2022B004 to X.C.).

